# Sexually Transmitted Infections Select for Different Levels of Immunocompetence And Reproductive Efforts in The Two Sexes

**DOI:** 10.1101/2021.01.07.425749

**Authors:** Lafi Aldakak, Frank Rühli, Nicole Bender

## Abstract

Sex differences in immunity have been described in humans and other mammal species. Females have a lower incidence of infections and non-reproductive malignancies and exhibit higher antibody levels after vaccination. Existing evolutionary explanations are based on differences in reproductive strategies and reaction to extrinsic differences in susceptibility and virulence between the sexes. Here, we test the hypothesis that known differences in the probability of transmission and outcome of sexually transmitted infections contribute to sex differences in immunocompetence. We modelled reproductive and immune investments against a fertility limiting Sexually Transmitted Infection (STI). We show that, in line with previous findings, increased susceptibility selects for tolerance to the parasite while increased virulence selects for resistance against it. Differences in reproductive strategies between the sexes lead to sex differences in immunocompetence, mostly with higher competence in females. Extrinsic differences in susceptibility and virulence between the sexes can augment or alleviate the evolutionary consequences of intrinsic differences depending on their direction and magnitude. This indicates that the selection of sex-specific immune strategies is less predictable than thought before and explains why sex differences in immunity have been found to be not universal and pervasive across animal species.

## 2. Introduction

Sex differences in immunity have been long observed in humans and other mammals (Schalk and Forbes 1997, Klein and Flanagan 2016). It has almost become common wisdom to consider males as the „sicker sex” (Zuk 2009, Bacelar, White et al. 2011). Many observations have contributed to forming this opinion. Females have a lower incidence of infections, develop higher antibody levels after vaccination, shed fewer viral particles, and suffer lower case fatality of many viral infections (Klein, Bird et al. 2001, Lin, Yao et al. 2006, Klein and Flanagan 2016). Even though these differences speak for a superior immune response in females, such generalisations may hide qualitative differences in the immune response between the sexes. The incidence of autoimmune diseases is strikingly higher in females, with 80% of autoimmune disease incidence occurring in women in humans (Cooper and Stroehla 2003, Cooper, Bynum et al. 2009). On the other hand, phenomena related to high immunopathology in the course of an infection, like the cytokine storm, are more common in males (Tisoncik, Korth et al. 2012). These observations indicate a more nuanced difference in the regulation of immune responses rather than an inferior male immune system.

The proximate reasons behind the difference in immunity have been well studied. Testosterone has been held culprit for the higher susceptibility to infections in males while oestrogen has been shown to enhance humeral immune responses (Klein and Flanagan 2016). The extra X-chromosome in females has also been associated with a bigger repertoire of adaptive immune components in females (Klein, Jedlicka et al. 2010, vom Steeg and Klein 2016). Although the universality of sex differences in immunity across animal species has been challenged recently, the aforementioned immune regulation differences in humans and mammals still require evolutionary explanations (Kelly, Stoehr et al. 2018). The first attempts to explain the sex difference in immunity from an evolutionary life-history perspective argued that the tradeoff between costly immune defences and reproductive efforts lead to males investing less in immunity to increase their mating success (Zuk 1990, Rolff 2002, Zuk and Stoehr 2002). According to this argument, stronger sexual selection on males should correlate with superior immune responses in females. Stoehr and Kokko (2006) tested this hypothesis in a modelling paper and showed that, while stronger sexual selection may decrease optimal allocation to immunity in males, males may invest more than females in immune defences if their attractiveness depends on their immunocompetence (Stoehr and Kokko 2006). Restif and Amos (2009) argued that Stoehr and Kokko’s model lacked a genetic framework, ecological and evolutionary dynamics to arrive at these conclusions. Given that ecological dynamics are crucial to determining the adaptive value of immunity in host-parasite interactions (Roy and Kirchner 2000, Restif and Koella 2003, Boots, Best et al. 2009, Donnelly, White et al. 2015), they revisited the general question asked by Stoehr and Kokko (2006) while incorporating ecological feedbacks between immune strategies and parasite prevalence and found that different reproductive strategies following Bateman’s principle alone are not enough to select for different levels of allocation to immunity between the sexes (Restif and Amos 2010). However, they found that females and males respond differently to extrinsic differences in the susceptibility to disease and virulence of parasite. For example, they predicted that higher exposure to infections in males selects for lower immunocompetence than in females. These results highlighted the importance of epidemiological feedbacks in studying the evolution of sex differences in immunity. Bacelar et al. (2011) found that male-biased parasitism evolves when males are subject to greater competition for resources or have a shorter lifespan, also highlighting the importance of accounting for ecological feedbacks (Bacelar, White et al. 2011). More recent attempts at explaining sex differences in immunity have considered tradeoffs around the magnitude and sensitivity of immune responses (Metcalf and Graham 2018). Surprisingly, modelling immune tradeoffs alone resulted in males aligning with higher immune sensitivity and females evolving greater magnitude of immune responses in contradiction with empirical observations. Only by assuming a higher infection risk and mortality around reproduction time in the sex that makes bigger parental investment (females) did their predictions match empirical observations with females evolving higher immune sensitivity. Although it did not account for ecological feedbacks between infection risk and immune strategies, Metcalf et al. (2018) is the first attempt to model sexual dimorphism around the sensitivity of immune surveillance and immunopathology, independent of tradeoffs around resource allocation to immunity.

We propose the hypothesis here that differences in susceptibility and outcome of Sexually Transmitted Infections (STIs) between the sexes select for different levels of immunity. STIs are particularly relevant to the evolution of sex-specific strategies and can practice selective pressures different from non-sexually transmitted diseases (Kokko, Ranta et al. 2002, Ryder, Webberley et al. 2005, McLeod and Day 2014). Many STIs are transmitted more efficiently from males to females. For example, the risk of genital herpes transmission in humans from a male to his female partner is 19%, whereas it is about 5% for transmission from a female to her male partner (Mertz, Benedetti et al. 1992). After a single episode of sexual intercourse, a woman has a 60% to 90% chance of contracting gonorrhoea from her infected male partner, whereas the risk of contracting gonorrhoea for a man is 20% to 30% when the female partner is infected (Hooper, Reynolds et al. 1978, Platt, Rice et al. 1983). The reasons for this difference include greater exposure in females as a result of pooled semen in the vagina and greater trauma to the surfaces during intercourse. Most cases of tubal factor infertility in women are attributable to untreated sexually transmitted diseases that ascend along the reproductive tract and are capable of causing tubal inflammation, damage, and scarring (Tsevat, Wiesenfeld et al. 2017). In men, semen quality deteriorates with STIs (Ochsendorf 2008). In non-human species, surveying two wild populations of stickleback confirmed that the presence of fibrosis scar tissue is associated with reduced parasite burden in both male and female fish. However, fibrotic fish had lower reproductive success (reduced male nesting and female egg), indicating big costs of the lingering immunopathology (De Lisle and Bolnick 2020).

But can STIs contribute to the evolution of sex-specific defence strategies?

Early models on STIs showed that the increased risk of catching the disease when mating could select females to decrease multiple mating (Thrall, Antonovics et al. 1997). Wardlaw et al. (2019) distinguished viability reducing STIs from fertility reducing STIs and showed that viability reducing STIs escalate sexual conflict while fertility reducing STIs de-escalate sexual conflict (Wardlaw and Agrawal 2019). Interestingly, McLeod et al. showed that it is advantageous for a sexually transmitted agent to be fertility limiting rather than viability limiting (McLeod and Day 2019). Johns and Henshaw et al. (2019) argued that males can benefit from transmitting sterilising STIs to their partners if STI-infected females invest more in their current offspring as a response to the reduced future reproductive prospects. (Johns, Henshaw et al. 2019). This can select males to evolve lower immunocompetence to acquire the STI and infect their mates (Lena, Pourbohloul et al. 2005).

Here, we use compartmental models of disease transmission to test the hypothesis that differences in susceptibility and outcomes of STIs contribute to the evolution of different immune strategies in the two sexes.

## 3. Methods

### 3.1. the epidemiological model

As demonstrated in figure 1, the basis of our model is a compartmental model of disease transmission that accounts for the feedback between disease prevalence and immune strategies. We assume that all individuals enter the model upon maturity in the susceptible compartments. We use the epidemiological model to derive the stable demographic structure of the population, assuming that demographic processes happen at much faster rate than evolutionary processes. We assume that the rates of STI transmission and mortality rates are fixed for each sex and that the STI is so cryptic that mate choice cannot favour mating with uninfected individuals. Assuming that *B* is the rate of recruitment of new virgin sexually mature individuals to the population (with equal newborn sex ratios), *β*_f_ is the rate at which females get infected (depends on the proportion of infected males in the population as explained below), *γ*_f_ is the rate at which infected females recover and *μ*_f_ is the natural mortality rate in females, the total change in the proportion of susceptible females is:

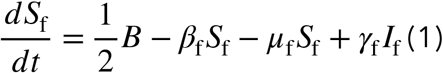

**Figure 1.**
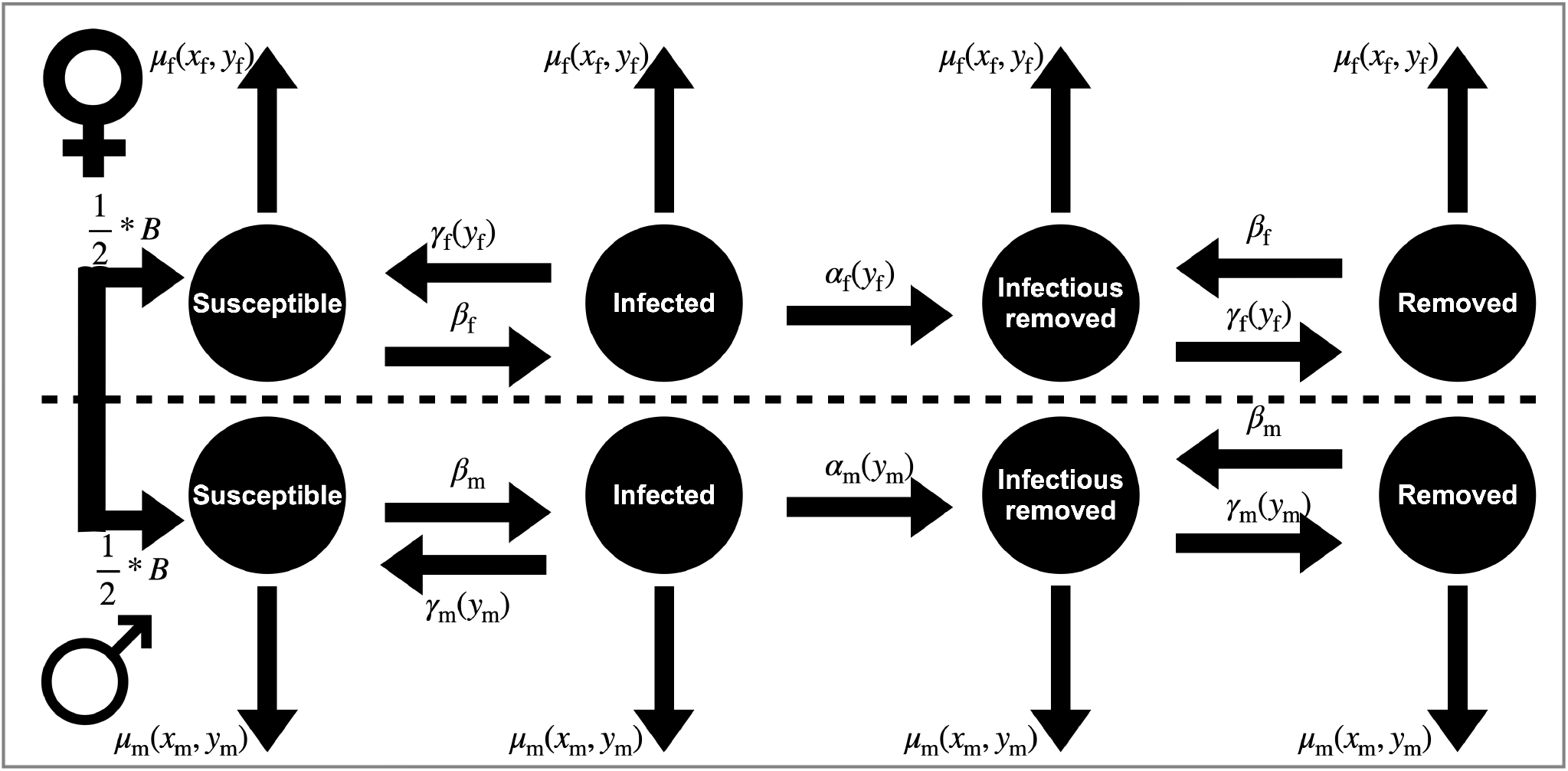
Schematic representation of the model. Females and males enter the population in the susceptible compartments. Upon mating with an infected male, a female becomes infected with a probability. Once infected, the female may recover at a rate, function of immune sensitivity or become sterile at a rate, sum of direct damage by the parasite and immunopathology. Infertile individuals may still mate and transmit the infection or recover and remain infertile. Males follow an analogous path. All individuals suffer a natural mortality rate, that is a function of both the mating investment and the immune sensitivity.

Similarly, infected females either recover at rate *γ*_f_, become sterile at rate *α*_f_ or die at rate *μ*_f_; therefore, the total change in the proportion of infected females is:

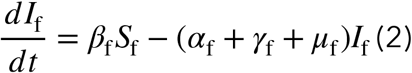

If a female becomes sterile during infection, she moves into the infected removed chamber (*IR*_f_). Sterile infected females do not go back to reproduction but they can still mate and spread the infection or recover and remain in the population until they die naturally. Consequently, we write the equation for change in the proportions of infected removed females:

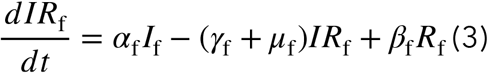

Lastly, infected removed females move into the removed chamber (*R*_f_) upon recovery. Removed females remain in the population until they die naturally or get reinfected upon mating with an infected male and become infected removed again, consequently:

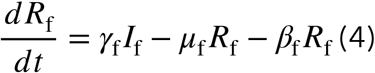

Similarly, the rates of change in males in each class are given by:

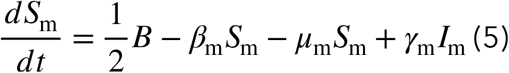

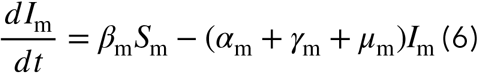

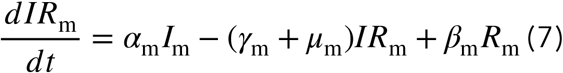

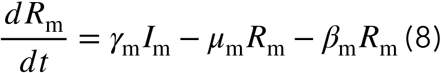

We assume that the population size has reached a stable equilibrium meaning that new individuals enter the population at the same rate as older individuals die. Therefore, the recruitment *B* is given by:

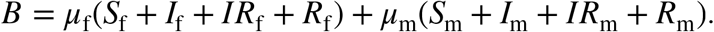

We can then set the derivatives in equations (1) to (8) to zero and solve numerically, under the constraint that *S*_f_ + *I*_f_ + *IR*_f_ + *R*_f_ + *S*_m_ + *I*_m_ + *IR*_m_ + *R*_m_ = 1 to find the composition of the population at equilibrium.

### 3.2. Immune sensitivity, reproductive efforts and mating rates

We assume that each individual has a set of two traits that determine its immune competence, mating rates, and, in one part of the model, natural mortality (explained below). Females allocate a proportion 0 < *x*_f_ < 1 to reproduction and have an immune sensitivity 0 < *y*_f_ < 1. The survival chances of a brood of offspring is proportional to the female investment *x*_f_ (for example when the number of offspring in one brood or their size depends on the females reproductive investment). Females mate upon reaching maturity and re-mate once their offspring reach maturity; therefore, female mating rates are independent of their investment strategies (i.e., there is never a shortage of males in the population). Assuming that the development time of offspring takes one time unit and that females suffer a constant mortality rate of *μ*_f_, she survives until her offspring mature with a probability 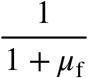. Therefore, the fitness gain for a female from a single mating equals the probability of surviving to produce offspring times her reproductive efforts: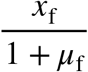

Similarly, males allocate 0 < *x*_m_ < 1 to reproduction and have an immune sensitivity of 0 < *y*_m_ < 1. When all males in the population have the same reproduction strategy *x*_m_, the share of matings by a male of a given infection status depends on the frequency of these males. The probability that any given mating is with a focal male of status *M* (where *M* = *S*_m_, *I*_m_, *IR*_m_ *or R*_m_) in this case is given by:

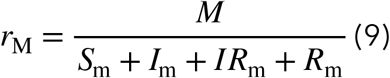

Similarly, the rate at which a male of infection status *M* mates with females of status *F* (where *F* = *S*_f_, *I*_f_, *IR*_f_ or *R*_f_) is:

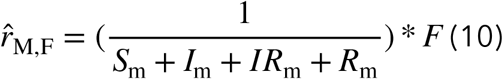

Meaning that 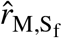 is the rate of mating with susceptible females, 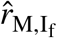 the rate of mating with infected females, 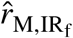 the rate of mating with infected sterile females and 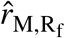 the rate of mating with uninfected sterile females. Note that the total mating rate of type *M* males with type *F* females equals that of type *F* females with type *M* males 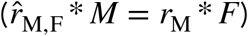, fulfilling the Fischer condition. The dependency of the mating success of a focal male on his mating investment is explained in the adaptive dynamics section.

Given that 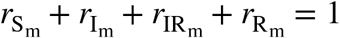, it follows that the probability that any given mating is with an infected male equals 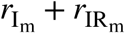 and the probability that a female becomes infected from any given mating is:

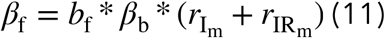

Following the same reasoning, the rate at which an uninfected male becomes infected depends on the rate at which he mates with infected females so that:

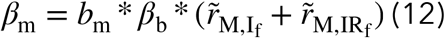

Where the parameter *β*_b_ determines the basal probability of infection when an infectious individuals mates with a susceptible one. The scaling factors *b*_f_, *b*_m_ allow as to account for biological structural sex differences in the susceptibility to infection upon an encounter with an infective partner. For example, *b*_f_ : *b*_m_ = 1 : 1 implies that the probability of transmission is equal for the two sexes. If *b*_f_ : *b*_m_ = 2 : 1 transmission is twice as likely if the infected partner is male.

### 3.3. The immune recovery-sterility tradeoff

Now we consider how recovery and sterility rates are determined by sex specific immune strategies. As explained above 0 < *y*_f_ < 1 is the sensitivity of the immune response in females and 0 < *y*_m_ < 1 is the sensitivity of immune responses in males. The recovery rates in females and males are determined by:

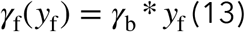

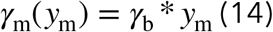

Where *γ*_b_ is a scaling factor. As sex-specific immune sensitivity increases, the corresponding recovery rate increases accordingly. The disadvantage of having highly sensitive immune responses is a higher level of immunopathology. High immunopathology increases the rate at which infected individuals become sterile due to tissue scarring and over-inflammation. Therefore, the sterility rate is the sum of direct damage caused by the parasite and collateral damage due to immunopathology so that:

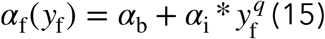

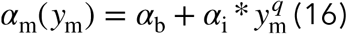

Where *α*_b_ is the intrinsic virulence of the parasite corresponding to the direct damage it does to reproductive tissues in order for it to replicate, *q* ≥ 1 is a parameter that determines the strength of the tradeoff between parasite clearing and collateral damage and corresponds to the overlap between parasite and host molecular signatures and the specificity of immune defences. The bigger *q* is, the weaker is the tradeoff between recovery and immunopathology.

The tradeoff between recovery rates and sterility rates around the parameter allows studying selection on immune sensitivity where host fitness may increase by increasing recovery rate (a type of resistance against the parasite) or decreasing the rate of sterility (a type of parasite tolerance corresponding to small values of *y* as described in (Raberg, Graham et al. 2009)).

### 3.4. Tradeoffs around resource and reproductive efforts

We use the framework presented above to study three scenarios distinguishing between selection due to the recovery-immunopathology tradeoff alone or in combination with tradeoffs around resource allocation to immunity and reproductive efforts. Hence, we construct the following three sub-models as follows:

(I) Immune sensitivity affects recovery and sterility rates as explained above. Natural mortality rate *μ* is a fixed parameter of the model. Hosts suffer costs of high immune sensitivity (higher rate of becoming sterile in the course of an infection) only when infected; therefore, costs of immunity are facultative.
(II) Immune sensitivity affects recovery and sterility rates as well as the natural mortality rates. As a result natural mortality rate is a function of sex specific immune sensitivity as follows:

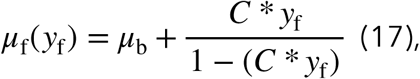

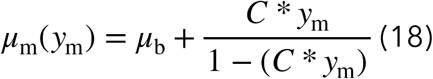 Natural mortality, as a result, is the sum of extrinsic mortality *μ*_b_ and an extra term that corresponds to the tradeoff between resources spent on immunity (*C* * *y*) and resources available for other types of somatic maintenance (1 – *C* * *y*). When immune sensitivity equals zero, the host suffer no extra mortality and its lifespan depends on extrinsic mortality alone. The more sensitive a host is and the higher the costs of an immune response are, the less left resources it has for somatic maintenance and the higher its natural mortality rate is.
(III) Immune sensitivity affects recovery and sterility rates as well as the natural mortality rates. Natural mortality depends on sex-specific immune sensitivity and resources invested in reproduction. As a result, natural mortality can be written as:

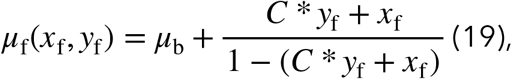

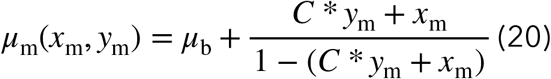 Meaning that mortality is an accelerating function of the total investment in immunity and reproduction. *x* + *y* = 1 implies instant death and 1 − (*x* + *y*) can be seen as the resources invested in somatic maintenance.

### 3.5. Expressions for fitness and Evolutionary Stable Strategies

Using the models presented above, we can derive the demographic make-up of the population at equilibrium when all individuals of each sex play the same strategies. Assuming that demographic change happens much faster than evolutionary change, we can study the conditions for the invasion of a rare mutant host into a resident population at its demographic equilibrium. We derive an expression for invasion fitness of a rare mutant across its life assuming the mutant is rare enough that we can ignore its effects on epidemiological dynamics. Upon sexual maturation, a mutant female with strategies 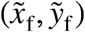 enters the population in the susceptible class where it mates with fertile males with a probability 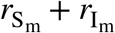. The fitness gain from any given mating with a fertile male is 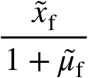, where the tilde denotes a parameter that may differ between the mutant and the resident population. The female is expected to remain in the susceptible class for 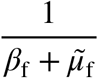. The mutant female become infected rather than dies with probability 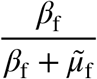. It remains in the infected class for a period 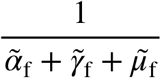 where it mates with fertile males with probability 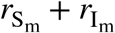 and gets fitness gains 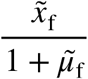 from each mating. The infected female may either become sterile due to infection and immunopathology, die naturally or recover with probability 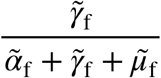 and return to the susceptible class to begin the cycle anew. We can, therefore, write the expression for the total reproductive output of a mutant female as the following recursion equation:

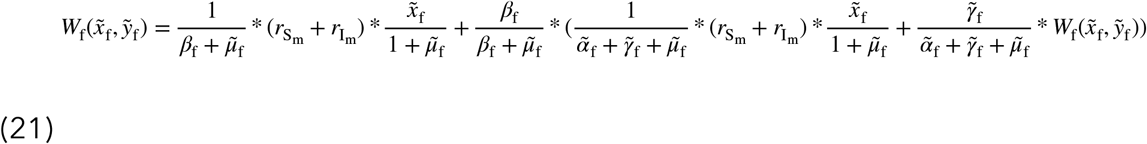

We can solve this for 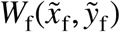, the fitness of a mutant female playing strategies 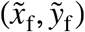.

The mating success of mutant male with reproductive investment 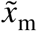 in a stable resident populations where males have reproductive investment *x*_m_ equals:

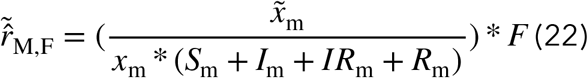

Using the same reasoning, we can write the expression for the invasion fitness of a mutant male playing 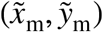 in a resident population where resident males play (*x*_m_, *y*_m_) strategies and resident females play (*x*_f_, *y*_f_) as follows:

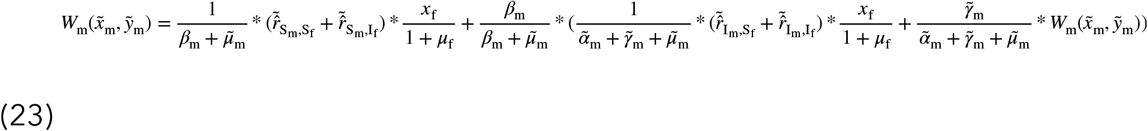

Which can also be solved for 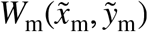. Relying on the simplifying assumption that the additive genetic variance is approximately equal for all traits, selection differentials for females and males are, as in (Johns, Henshaw et al. 2019), approximately proportional to:

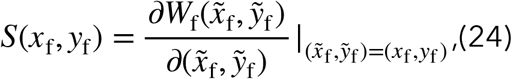

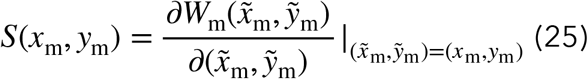

By starting with arbitrary initial strategies and following the selection trajectories defined by the selection differentials until converging to an equilibrium, we can find evolutionarily equilibria by iterating the equation with a small positive constant Δ:

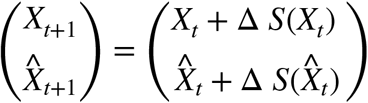

Where *X* is a vector of female strategies *X* = (*x*_f_, *y*_f_) and 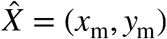 is a vector of female strategies. This means that the strategies move a small step in the direction of selection with each time step. The size of the step equals Δ times the selection intensity (Δ=0.01 was found to be suitable). Equations (1) to (8) were used to find stable population structure for each iteration.

## 4. Results

### 4.1. No extrinsic sex differences

#### (A) No reproductive allocation tradeoff, with or without energetic costs

If costs of high immune sensitivity are immunopathological only (model (I)) or immunopathological and energetic (model (II)), identical ESSs evolve in the two sexes (figure 2). When the susceptibility to the infection in both sexes increase simultaneously (*b*_f_, *b*_m_), lower immune sensitivity ESSs evolve. This means that higher transmission rates in these tradeoff structures favour tolerance over resistance, in agreement with previous theoretical results (Restif and Koella 2004, Restif and Amos 2010). Higher transmission rates make the infection more likely to occur and re-occur, leading to chronic infections. Therefore, the relative benefit of rapid recovery decreases and the effective costs of immunopathology increase, selecting for tolerance.

**Figure 2.**
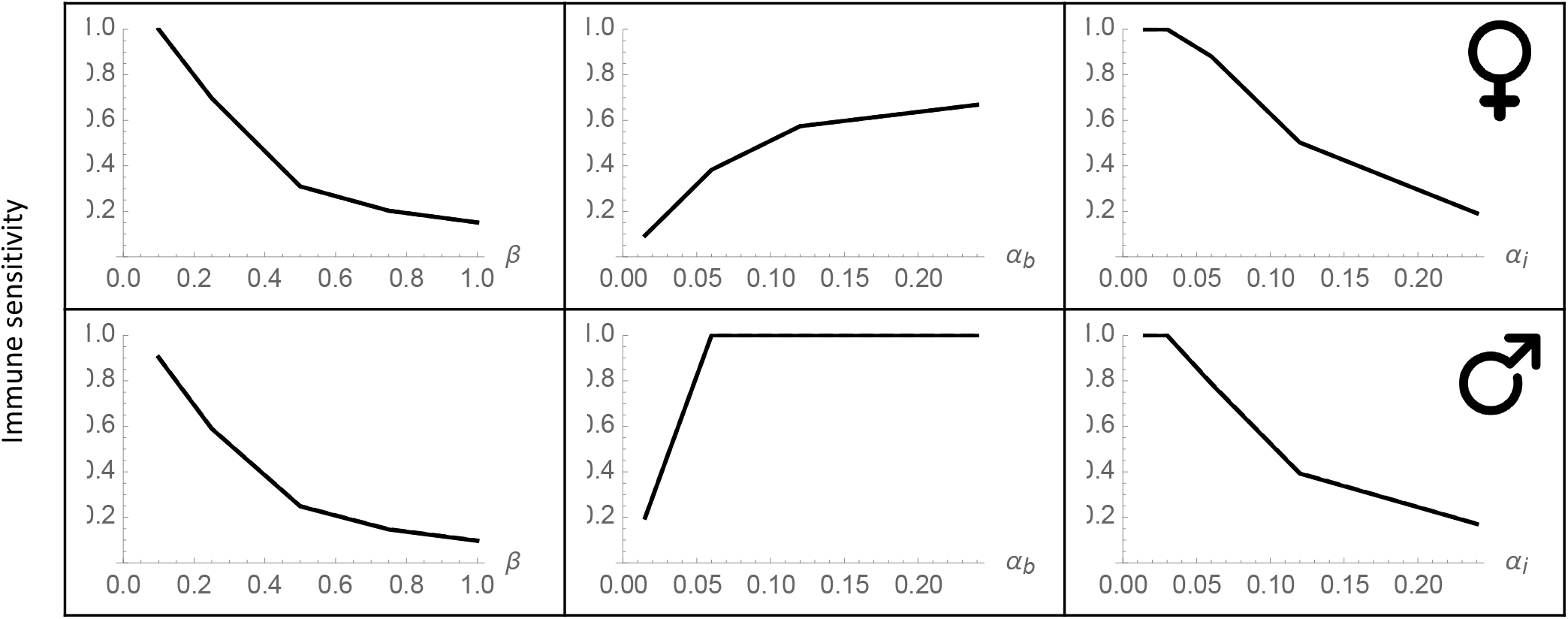
Immune sensitivity ESS against increasing susceptibility (left), virulence (middle) and immunopathology (right), in females (top) and males (bottom). The two sexes have similar values of immune sensitivity. Increased susceptibility and immunopathology favour lower levels of sensitivity while increasing virulence selects for higher sensitivity. Other parameter values: ***C* = 0.005, *μ*_b_ = 0.001, *q* = 2**

Increasing the direct damage caused by the parasite (*α*_b_) in both sexes simultaneously results in a monotonic increase in ESS immune sensitivity. Unlike non-sexually transmitted diseases, increased parasite virulence does not result in lower prevalence of the infection (infected hosts may become sterile but remain capable of transmitting the infection). Moreover, increasing direct parasite damage makes the infection more costly to individual hosts without affecting susceptible or recovered hosts; therefore, there is no incentive to evolve lower sensitivity as *α*_b_ increases.

Increasing the level of immunopathology associated with increased immune sensitivity (*α*_i_), selects for lower immune sensitivity ESS. This is biologically intuitive as increasing the costs associated with indirect damage by the immune response in comparison to the direct damage from the parasite increases the costs of high sensitivity and selects for, all else being equal, immune tolerance instead of resistance.

#### (B) Energetic costs and reproduction allocation tradeoffs

Now, we introduce resource allocation to reproduction (*x*_f_, *x*_m_) as a second evolving trait that can follow different trajectories in the two sexes (model III). As shown in figure 3, the two sexes evolve different immune sensitivity and allocation to reproduction ESSs. However, females and males do not exhibit qualitatively different responses to changes in susceptibility, parasite damage, immunopathology magnitude, or basic recovery rates to models I and II. In general, females exhibit larger immune sensitivity than males while males invest more resources in reproduction as females. The only exception to this pattern is in response to increased direct parasite damage *α*_b_ where male immune ESS exhibits slightly larger sensitivity to changes in direct damage than females. As *α*_b_ bincreases, immune sensitivity ESS increases slightly more steeply in males so that males have higher immune sensitivity against high values of *α*_b_. Allocation to reproduction exhibits the reverse pattern as female investment in reproduction exceeds male investment in response to increased parasite damage for the following reason. The increased parasite direct damage and the corresponding increased level of immune sensitivity increase the probability of sterility upon infection (the sum of direct and indirect damage). This limits the future reproductive perspectives of both females and males. Despite spending more resources on immunity, the consequently shorter fertile period creates more incentive to invest in reproduction, apparently so in males than in females, in a pattern similar to terminal investment.

**Figure 3.**
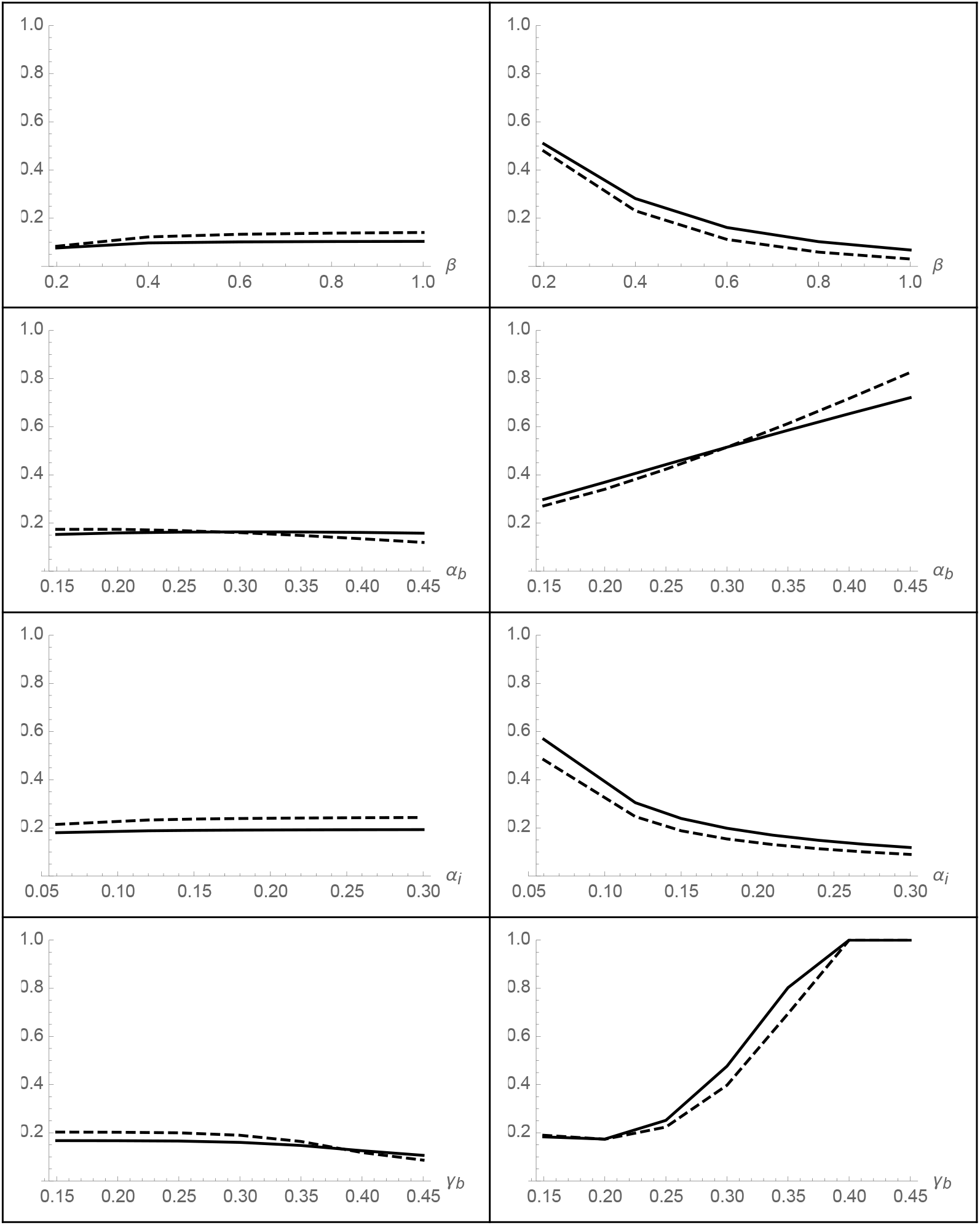
Optimal reproductive investments (left) and immune sensitivity (right) against increasing (from top to bottom): susceptibility, virulence, immunopathology and recovery. Solid lines resemble strategies in females while dashed lines resemble strategies in males. The response does not qualitatively differ between the sexes but females mostly have higher immune sensitivity. Other parameter values: ***C* = 0.005, *μ*_b_ = 0.001, *q*= 2**

**Figure 4.**
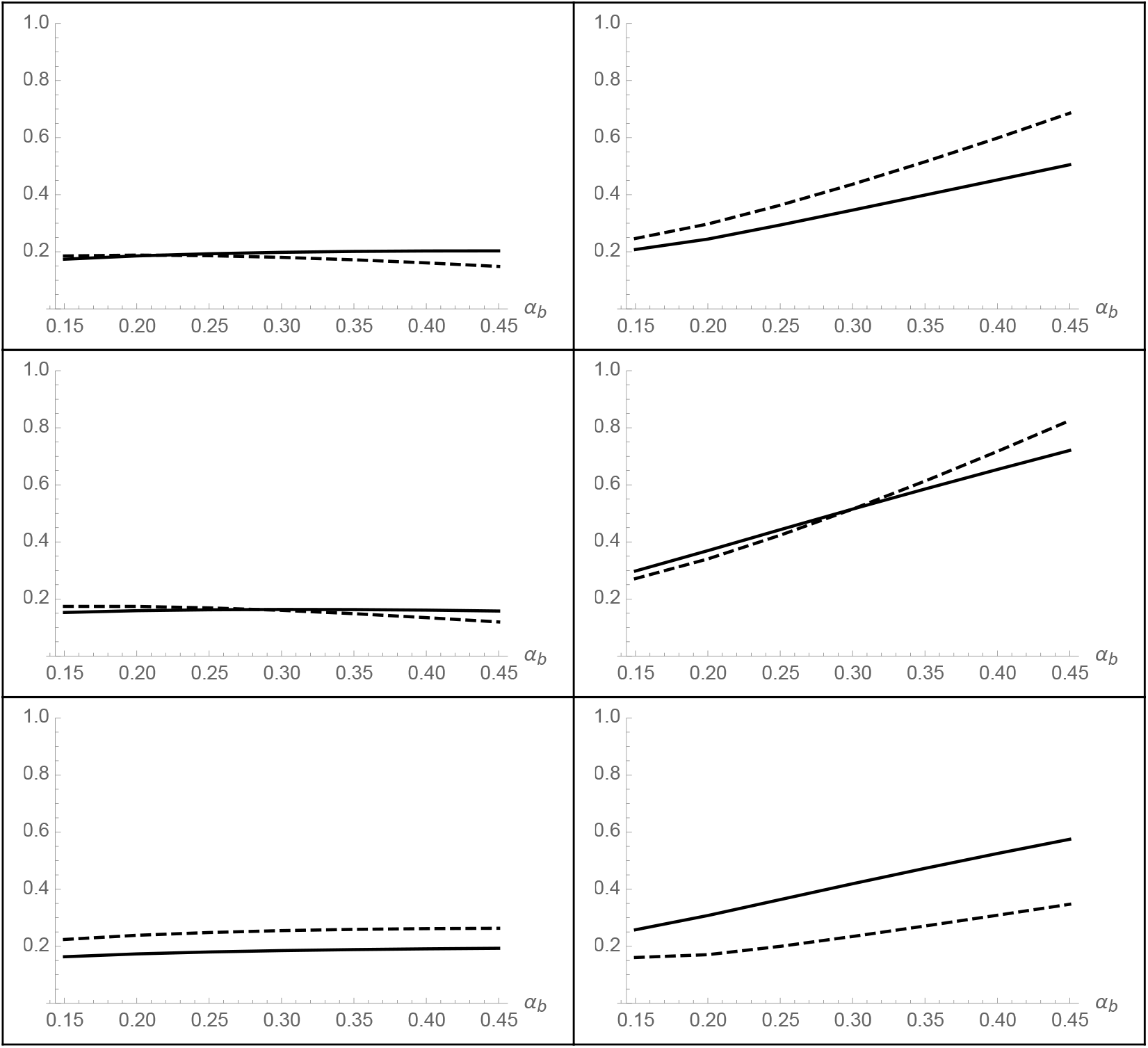
Optimal reproductive investments (left) and immune sensitivity (right) against increasing virulence. Solid lines resemble strategies in females while dashed lines resemble strategies in males. In the top row, females are two times as susceptible to the infection as males (***b*_f_ : *b*_m_ = 2 : 1**), the two sexes are equally susceptible in the middle row(***b*_f_ : *b*_m_ = 1 : 1**), and males are twice susceptible in the bottom(***b*_f_ : *b*_m_ = 1 : 2**),. Other parameter values: ***C* = 0.005, *μ*_b_ = 0.001, *q* = 2**

**Figure 5.**
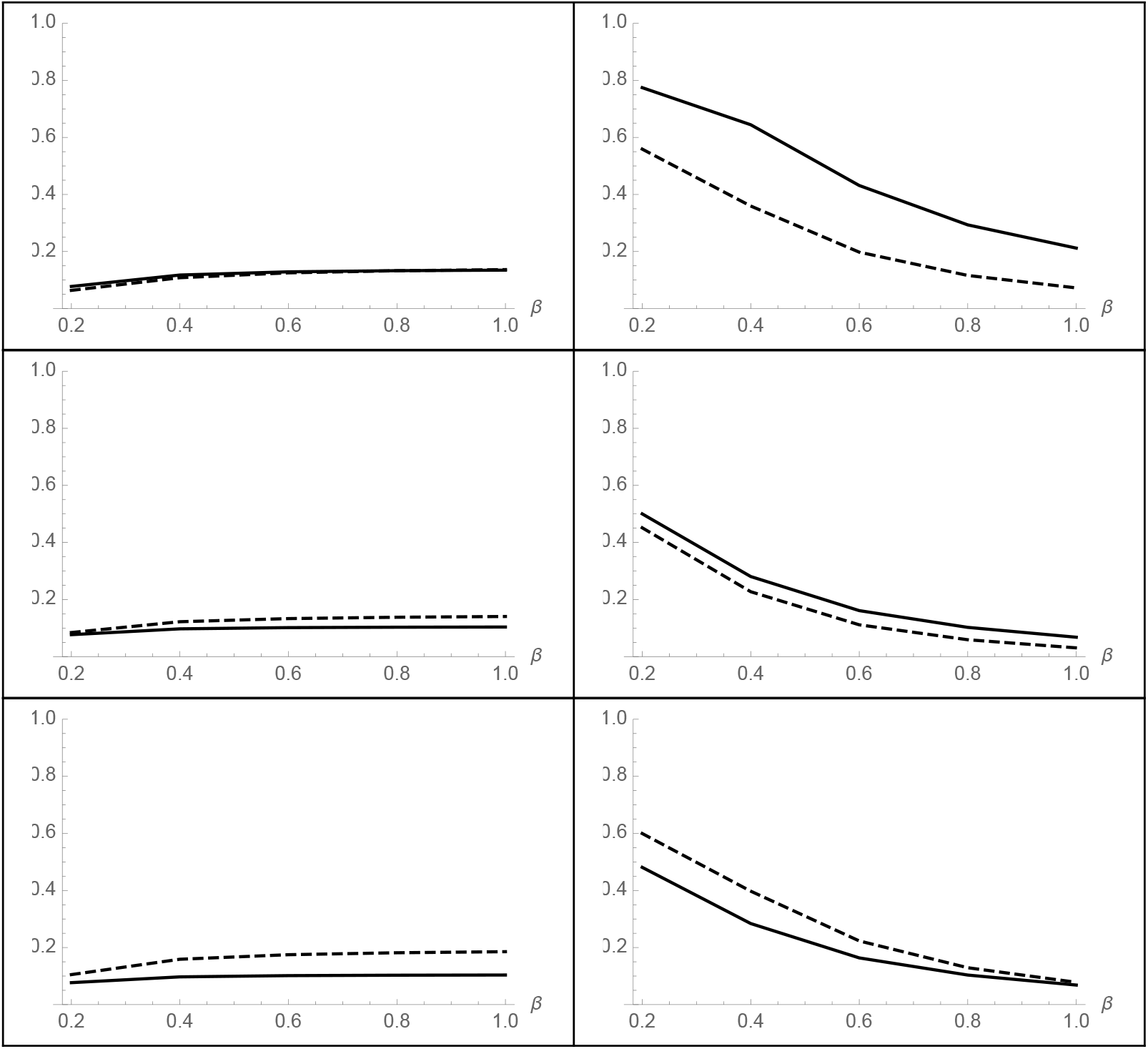
Optimal reproductive investments (left) and immune sensitivity (right) against increasing susceptibility. Solid lines resemble strategies in females while dashed lines resemble strategies in males. In the top row, parasite virulence is twice as much in females (***α*_f_ : *α*_m_ = 2 : 1**), the two sexes suffer equal virulence in the middle row(***α*_f_ : *α*_m_ = 1 : 1**), and malesahave twice the virulence compared to females in the bottom row(***α*_f_ : *α*_m_ = 1 : 2**),. Other parameter values: ***C* = 0.005, *μ*_b_ = 0.001, *q* = 2**

The only other case where females allocate more resources to reproduction than do males is against big values of the basic recovery rate *γ*_b_, in agreement with previous findings (Johns, Henshaw et al. 2019). Increasing *γ*_b_ corresponds to increasing the efficiency of the immune response in clearing out the infection before it leads to sterility. This selects both sexes to evolve higher immune sensitivity due to higher gains from increased sensitivity. Both sexes allocate less resources to reproduction, probably saving these resources for somatic maintenance. This can be understood by looking at equations (19) and (20) in model III. Allocating less resources to reproduction (smaller values of *x*_f_ and *x*_m_) decreases the second term in the natural mortality functions. This corresponds to investing less in current broods for females and mating competition for males to achieve gains in lifespan (thus increasing the accumulative number of broods for females or the reproductive period for males). This makes sense when the chances of sterility drop due to higher efficiency of immune defences and corresponds to terminal divestment (saving resources in current offspring to increase reproductive lifetime).

### 4.2. Extrinsic sex differences

Here, we study the evolution of immune sensitivity and allocation to reproduction (model III) when the two sexes differ in disease susceptibility beyond mating rates and reproduction patterns. This corresponds to anatomical or physiological differences that make the probability of transmission dependent on the sex of the infective partner. Namely, we allow the ratio between parameters *b*_f_, *b*_m_ to take different values to compare scenarios where the transmission likelihood is doubled from females to males (*b*_f_ : *b*_m_ = 2 : 1) or when transmission likelihood is doubled from males to females (*b*_f_ : *b*_m_ = 1 : 2) to the case where there is no sex difference in transmission likelihood. Analogously, we allow *α*_f_ : *α*_m_ to take values 2:1, 1:1, and 1:2 to test scenarios where the structure of the tradeoff between recovery and developing sterility is sex specific due to physiological and anatomical differences.

Changing the ratio of susceptibility to transmission does not change the general pattern in response to changes in other parameters. However, the more susceptible sex evolves lower immune sensitivity ESSs, in agreement with previous sections. For example, in response to increasing direct parasite damage, both sexes evolve monotonously higher resistance (higher immune sensitivity ESSs). As above, when females and males are equally susceptible to disease, females evolve higher ESS immune sensitivity in the lower range of while males evolve higher sensitivity in the upper range. When males are double susceptible, they evolve lower immune sensitivity ESSs even though they still respond to increasing parasite damage by increasing sensitivity. Therefore, the sex difference increases when males are more susceptible. When females are more susceptible to the infection, they evolve higher tolerance to the infection. This can reverse the general pattern and result in females evolving lower immune sensitivity and more tolerance across the whole range of parasite damage considered.

When the level of damage done by the parasite *α*_b_ differs between the sexes, higher resistance evolves in the sex that suffers greater damage. For example, males evolve lower immune sensitivity when *α*_b_ does not differ between the sexes. When parasite damage *α*_b_ is higher in females, the female-biased sex difference in immunity is bigger (higher sensitivity in females). When parasite damage is higher in males, however, males evolve higher levels of sensitivity than do females and this corresponds to a reverse in the general pattern and leads to higher tolerance to the parasite in females.

## 3.5. Discussion

Females and males can evolve different levels of allocation to reproduction and immune sensitivity in response to selection by STIs. Different mating efforts between the sexes (female investment affects the survival of the current brood while males compete over access to females) lead to different optimal strategies for immunity and reproduction in the sexes. Further differences in sex-specific susceptibility or virulence of the parasite can either augment the sex difference or alleviate it. We discuss the reasons behind these findings and compare them to previous results in the literature throughout.

In the absence of a tradeoff between reproductive efforts and immune competence, the two sexes evolve identical immune sensitivity ESSs. After including sex-specific tradeoffs between allocation to reproduction and remaining resources for immunity and somatic maintenance, the two sexes evolve sex-specific immune strategy, usually in the direction of higher resistance in females even in the absence of sexual selection and sex-specific parasitic impact. This contradicts previous findings by Stoehr and Kokko (2006) where the two sexes evolved identical strategies in the absence of sexual selection. The contradiction is a result of the difference in the structure of the tradeoff between their model and ours (Stoehr and Kokko 2006). In the aforementioned model, even though the male reproductive success was made dependent on the focal male trait relative to the rest of the population, in the absence of sexual selection, the structure of the function linking reproductive investment to fitness gains did not differ between the sexes. In the present model, female allocation to reproduction increases the survival of the current brood at the cost of the mothers` survival while males allocation increases their access to fertile females at the cost of their own survival. This structural difference leads to the two sexes aligning with different immune and reproduction allocation optima.

In their study about sex-specific immune defences against non-sexually transmitted diseases, Restif and Amos (2010) found that the intrinsic differences alone between the sexes are not enough to select for different levels of immunity (Restif and Amos 2010). Our results confirm their result that increased susceptibility selects for higher tolerance to parasites while increased virulence selects for resistance in both sexes. However, we show here that intrinsic sex-differences in mating efforts can lead to different immune strategies between the sexes. Furthermore, we show that extrinsic differences in susceptibility to or outcome of infections may either augment or alleviate existing sex-differences in immunity depending on whether they practice selection in the same or opposite direction as of existing intrinsic differences. Our results differ from those of Restif and Amos (2010) because we consider an additional tradeoff between immunity and allocation to reproduction. Even though Restif and Amos (2010) assume that males compete for access to females following Batemans (1948) principle, male reproductive success is not directly dependent on male strategies but rather proportional to its relative frequency and the function relating fecundity to immune strategy has the same shape in both sexes. We have shown here that males may evolve lower immune sensitivity and higher tolerance to parasites than females if their mating success depends on the resources dedicated to reproduction relative to other males in the population after allocating the rest to immunity and somatic maintenance.

We have shown that the response pattern to changes in disease susceptibility or consequences does not qualitatively differ between the sexes even though there might be quantitative differences in optimal sensitivity levels. Even though sex differences in variance in reproductive success (intrinsic differences) lead to generally higher sensitivity in females, ESS immune sensitivity in both sexes increases in response to increased parasitic damage and decreases in response to increased susceptibility (extrinsic differences). Because both intrinsic and extrinsic differences can affect the endpoints of selection on immunity, the effect of extrinsic differences can either mask or augment intrinsic sex differences in the immune response. For example, if mating differences lead to higher immune sensitivity in females in a given system, higher female susceptibility, which usually selects for tolerance, should select for lower sensitivity in females, dampening the sex difference in immunity. This might explain why a recent systematic review on the spread and magnitude of sex differences in immunity across animal species found them to be less universal than originally expected (Kelly, Stoehr et al. 2018).

Metcalf and Graham (2018) carried out one of the first attempts to employ tradeoffs between immune sensitivity and specificity to explain sex differences in immunity (Metcalf and Graham 2018). They verbally argued that „greater variability in male reproductive success means that males obtain greater fitness gains for investment toward securing mating opportunities than females“. They found that, assuming males should reduce investment in the magnitude of the immune response or the resolution of immune discrimination mechanisms, males evolve higher immune sensitivity than females. The reason behind the contradiction with our results is that this assumption overlooks the link between immune sensitivity and the accumulation of resource-costs of immunity. Increasing immune sensitivity lowers the threshold for triggering an immune response, increasing the overall number of immune reactions in the lifetime of the host and possibly even the magnitude of the response (higher mobilisation of immune effector cells and chemical mediators). Incorporating a direct tradeoff between immune sensitivity and resources left for reproduction and somatic maintenance in our model, we show that such verbal arguments can be misleading and males evolve lower immune sensitivity under a wide range of parameters under immune tradeoffs.

In a previous study on the evolution of immune sensitivity under immunopathology and autoimmunity costs, we have shown that high virulence is double problematic because it does not only imply high direct damage by the parasite but also selects the host to exhibit higher levels of immunopathology and low tolerance, increasing the fatality of the infection (Aldakak, Rühli et al. 2020). Analogously, we show here that in the context of STIs, high direct damage by the parasite selects for high immunopathology. This implies that STIs that cause a lot of direct damage to the host are similarly double problematic; they select the host to exhibit high levels of immune sensitivity accompanied by immunopathology. Echoing the effect of virulence in non sexually transmitted diseases, the resultant high probability of sterility after virulent STI infections resemble a combination of big direct damage by the parasite and the adaptive value of high resistance against such infections at the cost of high immunopathology.

Interestingly, females’ reproductive investment exceeds that of males in response to increased parasitic damage *α*_b_ and the accompanying adaptive high immunopathology (figure 3). This echoes results from Johns et al. (2019) that females infected with sterility-inducing STIs may increase their investment in their current brood due to diminished future reproduction (Johns, Henshaw et al. 2019). We show here that the likelihood of this terminal investment is higher the more virulent the parasite causing the STIs is. Also in agreement with Johns et al. (2019), when the efficiency of immune responses is high (big *γ*_b_), both sexes decrease their current reproductive allocation due to diminishing probability of sterility and bigger gains from investing in immunity.

We have not directly considered sexual selection in this study but we predict that strong sexual selection can inflate immune sex-differences when females are the choosey sex and males do not obtain attractiveness gains from higher immunocompetence. Once we release these assumptions, the pattern gets more complicated (in a similar way to Stoehr and Kokko (2006)) and should be addressed in a separate model. We have not considered the effects of immune memory in this model since repeated infections are common with STIs, decreasing the plausibility of immune memory (Leichliter, Ellen et al. 2007). Nonetheless, immune memory might theoretically change the predictions of evolutionary immunology due to its epidemiological effects on the prevalence of the infection (Boots, Donnelly et al. 2013). Moreover, trans-generational immune memory through antibodies transmitted from mothers to offspring may create additional fitness gains from immunocompetence in females leading to different sex-specific optima (Grindstaff, Brodie et al. 2003). Our study sheds light on the role of intrinsic and extrinsic differences in the susceptibility and outcome of STIs in the evolution of sex-different immune strategies and the shortcomings addressed here are an incentive to handle them more extensively in upcoming studies.

## Author contributions

L.A., N.B. and F.R. conceived the project. L.A. designed and performed the mathematical modeling and wrote the first draft; all authors revised and edited the manuscript.

## Acknowledgments

We thank Matthias Galipaud and Jonathan Henshaw for helpful comments that helped realise this project. The Mäxi Foundation, Zurich, funded the project through a grant to F.R.

